# Exposure to a hypercaloric diet produces long lasting changes in motivation^1^

**DOI:** 10.1101/2022.08.19.504605

**Authors:** Wendy Andrea Zepeda-Ruiz, Héctor Alan Abonza Paez, Marco Cerbon, David N. Velazquez Martinez

**Affiliations:** Departamento de Psicofisiología, Facultad de Psicología, Universidad Nacional Autónoma de México. Ciudad de México, México, 04510; Unidad de Investigación en Reproducción Humana, Instituto Nacional de Perinatología-Facultad de Química, Universidad Nacional Autónoma de México. Ciudad de México, México, 04510

**Keywords:** motivation, hypercaloric diets, breaking point, progressive ratio schedule, time on diet

## Abstract

Changes in motivation have been observed following induction of diet-induced obesity. However, to date, results have been contradictory, some authors reporting an increase in motivation to obtain palatable food, but others observing a decrease. Observed differences might be associated with the length of both the evaluation period and exposure to the diet. Therefore, the aim of this study was to evaluate changes in motivation during 20 weeks of exposure to a hypercaloric diet. Performance of the subjects in a progressive ratio schedule was evaluated before and during the exposure to a high-fat, high-sugar choice diet (HFHSc). A decrease in motivation was observed after 2 weeks of diet exposure, low levels of motivation remained throughout 20 weeks. A comparable decrease in motivation took longer (3 weeks) to develop using chow diet in the control group. Overall, our results suggest that, when changes in motivation are being evaluated, long periods of diet exposure made no further contribution, once motivation decreased, it remained low up to 18 weeks. Exposure to a HFHSc diet is a useful animal model of obesity, since it replicates some pathophysiological and psychological features of human obesity such as an increase in fasting glucose levels, body weight and the weight of adipose tissue.

## 1. Introduction

Obesity is a chronic, multifactorial disease (Ortiz & Kwo, 2015). Specifically, diet-induced obesity is the consequence of a sustained positive energy balance and is associated to pathophysiological (*e.g*., type 2 diabetes, hypertension, metabolic syndrome and hyperglycemia) (Dutheil et al., 2016; Ortiz & Kwo, 2015) and psychological (*e.g*., depression, impulsivity) (Sharma & Fulton, 2013; Smith & Smith, 2016) alterations.

To study psychological alterations related to obesity, the diet-induced obesity is the most widely used animal model. In this model, subjects have access to high-fat or high carbohydrates diet or to a cafeteria diet, where different food types are simultaneously presented (Lutz and Woods, 2012).

Changes in motivation have been reported in animal models of diet-induced obesity (Arcego et al., 2020; Fam et al., 2022; La Fleur et al., 2007). Motivation is defined as a process that allows organisms to approximate and avoid stimuli depending on hedonic or aversive characteristics (Salamone et al., 2016). It is worth noting that, according to the incentive salience model posited by Berridge (1996), it is possible to differentiate between two psychological components of motivation: “wanting” and “liking”; liking refers to the hedonic properties of food such as flavor or texture, while wanting refers to motivational aspects, specifically the incentive value associated to searching, approximation and consumption of food (Berridge et al., 2010).

Progressive ratio (PR) schedules have been widely used to evaluate the incentive value assigned to food. In these schedules, the response requirement to obtain a reinforcer increases in each trial or successive ratio, the most common parameter obtained in this schedule is the breaking point (BP), defined as the last completed ratio before rats stop responding for a criterion period established arbitrarily, e.g. five minutes (Valencia-Torres et al., 2014).

As previously mentioned, some authors have reported that exposure to an obesogenic diet produces motivational changes. However, some reported results are contradictory, subjects exposed to a palm oil saturated high-fat diet for 12 weeks achieved lower BPs in a PR schedule compared to a control group (Décarie-Spain et al., 2018); similar results were also found by Arcego et al. (2020) and Fam et al. (2022) who reported that after exposure to high-fat diet or a cafeteria diet, subjects showed a blunted motivation for sucrose.

On the other hand, La Fleur et al. (2007) and Sketriene et al. (2021), observed that subjects that had access to a high-fat, high-sugar choice diet (for 10 or 8 weeks, respectively) achieved higher BPs and increased the number of earned rewards. In addition, Figlewicz et al. (2014) also reported that after 3 weeks of a moderate high-fat diet exposure, subjects increased their active lever presses in comparison to subjects fed with a regular chow diet.

At least two factors may participate in the discordant motivational changes observed, these are the time of exposure to the diet and the type of diet (La Fleur et al., 2007; Figlewicz et al, 2014). Regarding the type of diet, high-fat diets are commonly employed because they produce a reliable increase of the subjects’ body weight. However, as Peris-Sampedro et al. (2019) suggested, it is also possible that food with a high content of carbohydrates could play a role in the psychological and physiological changes associated to obesity. In line with this observation, Gao et al. (2017) found that an inflammatory response was observed in the hypothalamus of animals fed with a high-fat diet with a high content of carbohydrates, but not in subjects exposed only to a high-fat, low content of carbohydrates diet. Therefore, the use of diets where both nutrients are available such as cafeteria diet or high-fat, high-sucrose choice (HFHSc) diet is desirable for the study of psychological and pathophysiological changes observed in obesity (Lutz & Woods, 2012).

Concerning the availability of the diet, Tracy et al. (2015) evaluated the performance of subjects in a PR schedule after different periods of diet availability and found that after 3 weeks, subjects on a chow diet and subjects on a high-fat diet achieved similar BPs; however; after 6 weeks, subjects on the high-fat diet achieved lower BPs, this suggests that if the period of diet exposure increases, changes in motivation are more dramatic.

As it has been suggested that a decrease in motivation depends on the time of diet exposure (Tracy et al. 2015) and the type of diet is also a relevant variable to consider when evaluating motivation, the present study sought to evaluate changes in motivation during 20 weeks of a HFHSc diet compared to access to a chow diet. The impact of the HFHSc diet on body weight, adipose tissue and glucose levels was also evaluated.

## 2. Materials and methods

### 2.1. Subjects

Twenty-four male Wistar rats (breeding colony of the Facultad de Psicología, Universidad Nacional Autónoma de México), 60 days old and weighing 250-300 g at the start of the study were individually housed in controlled conditions of temperature and humidity, under a 12:12 light-dark cycle. All rats had initial continuous access to tap water and a pelleted rodent diet (Rodent laboratory Chow 5001, PMI Nutrition International L.LC., Brentwood, USA). At their arrival to the lab, body weight was recorded for 5 consecutive days under free feeding conditions. Throughout the operant experiment, subjects were maintained at 85% of their initial free-feeding body weights with a correction for their natural growth. Tap water was freely available in their home cages. All procedures, housing and handling observed the National Institutes of Health guide for the care and use of Laboratory animals (NIH Publications 8^th^ Ed., 2011) and the study had the approval of the Ethics Committee of the Facultad de Psicología, Universidad Nacional Autónoma de México.

### 2.2. Apparatus

The rats were trained in operant conditioning chambers (Lafayette Instruments, Lafayette, IN, USA). One wall of the chamber had a recess into which a peristaltic bomb dispenser could deliver 0.2 ml of a 10% sucrose solution. A retractable lever inserted into the chamber through an aperture situated 8 cm above the floor and 5 cm to the right of the dispenser could be depressed by a force of approximately 0.2 N. The chambers were enclosed in a sound attenuating chest with rotary fans and masking noise. Experimental events and responses were controlled or recorded with a MED Associates interface (MED Associates, Inc. Fairfax, VT, USA) and a computer located in the same room.

### 2.3 Procedure

#### Behavioral training

One week before the start of the behavioral training, subjects were food deprived and their body weight gradually reduced to the 85% of their free-feeding period. Then, during two sessions, rats received 0.2 ml of a 10% sucrose solution under a fixed time schedule (FT20 sec), but a lever press was immediately followed by another 0.2 ml of sucrose; those rats that did not learn to press the lever were shaped manually. Thereafter, subjects were exposed to increasing fixed ratio schedules (1, 3, 5, 8 and 10) for 2 sessions each. After the last session of FR10 schedule, training in the PR schedule began. The PR schedule was based in the exponential progression: 1, 2, 4, 6, 9, 12, 15, 20…, derived from the formula (5xe^0.2n^)-5, rounded to the nearest integer, where *n* is the position in the sequence of the ratios (Bradshaw & Killeen, 2012; Roberts & Richardson, 1992). Experimental sessions took place once every day (between 11:00 and 13:00 h), in the light phase of the daily cycle, seven days per week and lasted 45 minutes. Subjects were trained until they achieved behavioral stability defined as less than 15% of variability in their breaking point (BP) during the last 10 sessions.

Subjects were exposed to the HFHSc diet for 20 weeks (see below) and the performance of the subjects in the PR schedule was evaluated (at least twice a week) from week 2 to week 20, we did not evaluate the performance on the first week to allow subjects familiarize with the diet.

#### Diet-induced obesity

Diet-induced obesity was carried out with a modified version to the protocol of Peris-Sampedro et al. (2019) of a high-fat, high-sugar choice diet (HFHSc). In short, subjects were assigned in a counterbalanced way to Control (C) and experimental group (E), according to the mean of the BP of the last 10 sessions of training. Subjects of both groups had *ad libitum* access to tap water and standard diet (5001, rodent laboratory chow, Purina^®^). Additionally, a HFHSc diet was offered to the E group, and it consisted of a 10% sucrose solution and a high-fat diet that consisted in INCA^®^ butter (ACH Foods México) that contains edible tallow and 0.004% of antioxidants. Body weight, water, standard diet, INCA^®^ butter, and sucrose intake were recorded daily, spillage on the bedding of the home cage was also recorded. Energy density of the food types were: 4.07 kilocalories (kcal)/g for the 5001, rodent laboratory chow, 9 kcal/g for INCA^®^ butter, and 4 kcal/g for the sucrose solution. Both groups were maintained in their diets for 20 weeks. In order to evaluate physiological changes associated to the HFHSc diet, at the end of the 20 weeks, subjects were food deprived for 6 h and the blood glucose from a tail-vein blood sample was quantified, according to the protocol of Furman (2015). The next day, subjects were sacrificed, and retroperitoneal, abdominal and perigonadal white adipose tissue was dissected and weighed.

### 2.4. Data Analysis

A two-way mixed ANOVA (groups x weeks) and pairwise comparisons with Bonferroni correction in the within-subjects factor were used to compare the weight and kilocalories intake of both groups and to compare the BPs before and after the exposure to a free-choice diet as well as the adipose tissue weight. When sphericity criterion was not met, Huynh-Feldt adjustment was used for the analysis of the main effects of the within-subjects factor. Glucose levels were compared with an independent sample t test. The significant level was set up to α=0.05.

## 3. Results

### 3.1. HFHSc diet produces an increase in kilocalories intake and body weight

After behavioral stability was achieved in the PR schedule (90 experimental sessions), subjects were exposed to their respective diets. After 11 weeks of access to the HFHSc diet, the total intake of kcal was larger in the E group in comparison to the C group (Two-way mixed ANOVA, F_(1,22)_= 22.926, p=0.01, η^2^_p_=0.510) (Fig. 1A). Pairwise comparisons in the within-subjects factor (weeks) (Two-way mixed ANOVA, Huynh-Feldt adjustment, F_(9.42,207.26)_= 21.993, p<0.05, η^2^_p_=0.500) revealed that in both C and E groups there was a decrease of their total kcal intake during week 20 compared to the first week (p=0.05 and p=0.05, respectively) (Fig. 1A).

**Figure 1.**
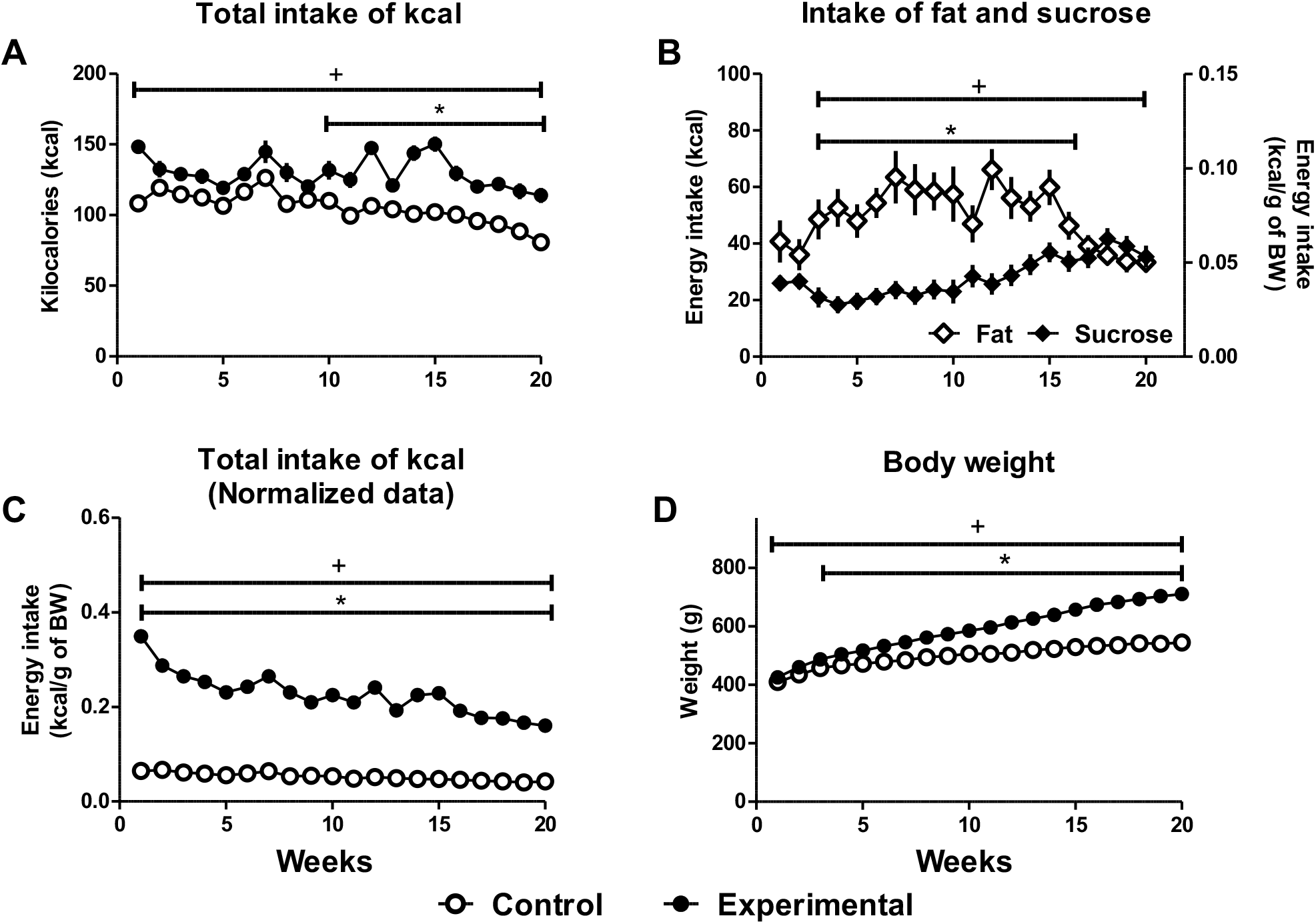
Intake and body weight on the HFHSc diet of the experimental and control group. A: Mean of total intake of kcal. B: Kcal intake from fat and sucrose of the experimental group. C: Normalized data (kcal/g of body weight) of total kcal intake. D: Mean body weight of subjects of control and experimental group during diet-induced obesity. Mean ± SEM. Horizontal bars with * show significant differences between groups (p<0.05) or nutrients; horizontal bars with + show significant differences between weeks (p<0.05).

After normalization of total kcal intake to body weight, we found that subjects from E group had a larger intake than the C group from the first week of access to the HFHSc diet (Two-way mixed ANOVA, F_(1,22)_= 1206.57, p=0.01, η^2^_p_=0.982). We also found that in both, C and E groups (Two-way mixed ANOVA, Huynh-Feldt adjustment, F_(7.80,171.68)_= 78.88, p<0.05, η^2^_p_=0.782) normalized total kcal intake was lower during week 20 compared with week 1 (p=0.05 and p=0.05, respectively) (Fig. 1C).

The last results may sound counterintuitive; however, it is necessary to consider that the elevated intake of kcal during the first week of the protocol was associated with the fact that subjects were food deprived during behavioral training on the PR schedule and then had *ad libitum* access to their diets, these procedures gave rise to an increase in their food intake that stabilized throughout the diet-induced obesity protocol.

Kcal intake from fat and sucrose was also analyzed. The kcal intake from fat was larger than for sucrose (Two-way ANOVA repeated measures, food factor, F_(1,11)_= 11.448, p=0.01, η^2^_p_=0.510) from week 3 up to week 15. After that, the kcal intake from sucrose increased, whereas the kcal intake from fat decreased (Two-Way ANOVA repeated measures, with Huynh-Feldt adjustment in the weeks factor F_(9.21,38.06)_= 4.965, p=0.01, η^2^_p_=0.311). In this way, rats adjusted their diet composition but their total kcal intake from both types of food was similar during the last 5 weeks (p=0.096) (Fig. 1B). When data of kcal intake from fat and sucrose was normalized by body weight, significant differences were still found (Two-way ANOVA repeated measures, food factor, F_(1,11)_= 746.44, p=0.01, η^2^_p_=0.985) and the same pattern of high fat intake followed by high sucrose intake described previously (Two-Way ANOVA repeated measures, with Huynh-Feldt adjustment, F_(8.66,57.40)_= 6.362, p=0.01, η^2^_p_=0.366) was found through the weeks (p=0.096) (Fig. 1B).

Body weight of both groups was also compared and we observed that from week 3, body weight was larger in the E group (Fig. 1C) (Two-way mixed ANOVA, F_(1,22)_= 27.097, p<0.05, η^2^_p_=0.552) compared with the C group. There were also, significant differences in the weeks factor (Twoway mixed ANOVA, with Huynh-Feldt adjustment, F_(1.71,38.31)_= 343.34, p=0.01, η^2^_p_=0.940). Although both groups increased their body weight throughout the manipulation (p=0.01), pairwise comparisons revealed that in the C group the body weight was similar between weeks 13 and 20 (p=1.0), while in the E group, there were significant differences in the body weight along the procedure, except between weeks 19 and 20 (p=0.223) (Fig. 1C).

### 3.2. HFHSc diet produces an increase in adipose tissue weight and elevated fasting glucose levels

The weight of peritoneal, retroperitoneal and gonadal adipose tissue of subjects was larger in the E group in comparison to the C group (Fig. 2A) (Two-way mixed ANOVA, F_(1,22_ = 75.206, p=0.01, η^2^_p_=0.774). There were also significant differences between the C and E groups when data from adipose tissue were normalized by body weight (tissue weight/g body weight) (Twoway mixed ANOVA, F_(1,22)_= 53.461, p=0.01, η^2^_p_=0.708). Glucose levels after 6 h of fasting were higher in the E group compared to the C group (Fig. 2B) (t_(22)_= −4.922, p=0.001).

**Figure 2.**
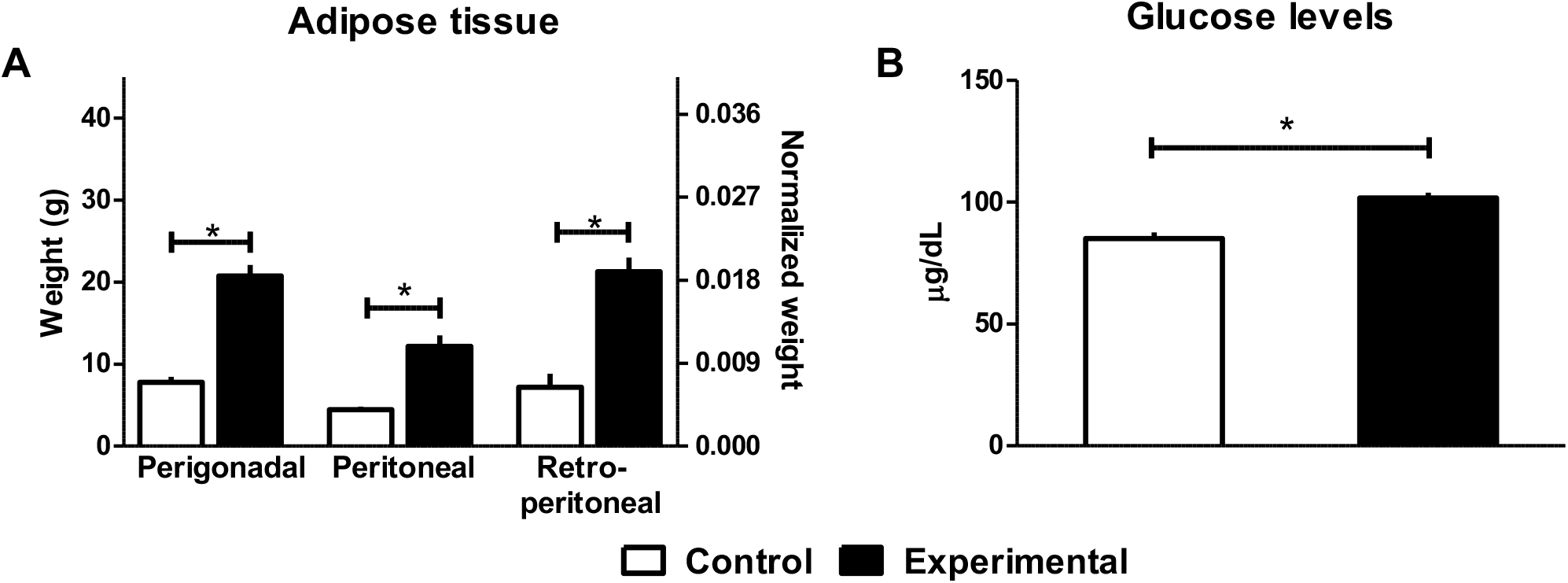
Adipose tissue weight and fasting glucose levels after exposure to the HFHSc diet. A: Perigonadal, peritoneal and retroperitoneal adipose tissue weight. B: Fasting glucose levels. Mean ± SEM. Horizontal bars with * show significant (p<0.05) differences between groups.

### 3.3. Breaking point after HFHSc diet was lower compared with that achieved during training

The BP achieved during the training phase was compared to the BP obtained after three weeks of the HFHSc diet protocol, we found that, in comparison with the training phase, subjects of C group achieved a lower BP after two weeks of standard diet, while the decrease of the BP in the E group occurred faster and was observed from the first week of access to the HFHsc diet (Fig. 3A) (Two-way mixed ANOVA, F_(2,44)_= 40.761, p=0.01). When the BP was achieved during the 20 weeks that were considered, we confirmed the decrease of the BP in the E group after one week of access to the hypercaloric diet and after two weeks in the C group. In addition, we observed that once the BP of C and E groups decreased it was maintained at a lower level until week 20 (Two-way mixed ANOVA, with Huyhn-Feldt adjustment; F_(5.59,123.17)_=26.967, p=0.01, η^2^_p_=0.551) (Fig. 3B).

**Figure 3.**
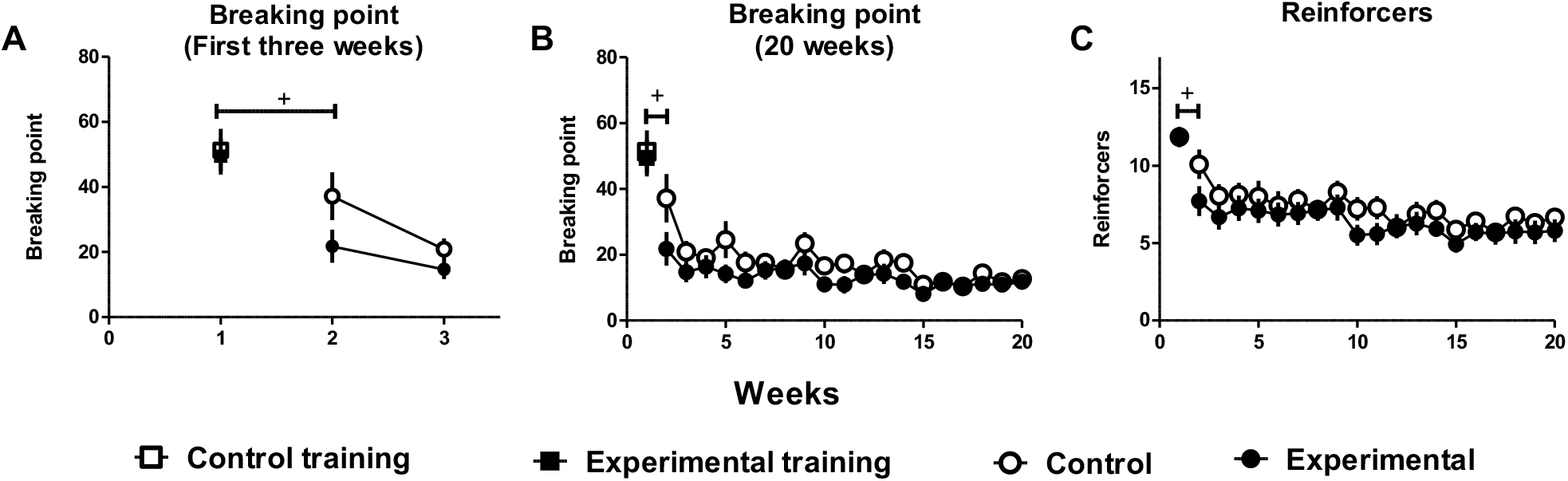
Breaking points achieved before (training) and during the HFHSc diet of the experimental and respective period of control group. A: Mean of the breaking point achieved during training and the three initial weeks of the HFHSc diet protocol. B: BPs achieved during the training phase and the 20 weeks of the HFHSc diet protocol. C: Earned reinforcers during the 20 weeks of access to the hypercaloric diet. Mean ± SEM Horizontal bars with + show significant differences between phases (p<0.05).

When the number of obtained reinforcers was compared, we also observed that both groups earned fewer reinforcers during the diet-induced obesity protocol (Two-way mixed ANOVA, with Huynh-Feldt adjustment, F_(9.57, 210.71)_=20.383, p=0.01, η^2^_p_=0.481) compared to the training phase; subjects of the C group earned fewer reinforcers after 3 weeks of standard diet (p=0.01), while the E group earned fewer reinforcers after 2 weeks of HFHSc diet (p=0.01). There were no significant differences between groups during training, nor during the diet-induced obesity protocol (Two-way mixed ANOVA, group factor; F_(1,22)_=1.129, p=0.300, η^2^_p_=0.049) (Fig. 3B).

## 4. Discussion

In the present study, we evaluated the performance of the subjects in the PR schedule during 20 weeks of exposure to the HFHSc diet. A decrease in BP during the access to the hypercaloric diet was accompanied by an increase in the body weight, in the adipose tissue weight and in the fasting glucose levels.

Although subjects of C and E groups increased their body weight during the exposure to the HFHSc diet it is worth noting that, in the E group, the increment in its body weight was observable during almost all the duration of the period of exposure compared to the C group, where subjects only increased their body weight up to week 13 and thereafter remained with a similar body weight until the end of the experiment. The increase in body weight of E group is in accordance to the results reported by Peris-Sampedro et al. (2019), who observed that even after three weeks on a HFHSc diet, subjects achieved a larger body weight in comparison with subjects fed with a standard chow diet.

We also evaluated other parameters that have been reported in animal models of diet-induced obesity, such as adipose tissue weight and fasting glucose levels (Ortiz and Kwo, 2015). An increase in perigonadal, peritoneal and retroperitoneal white adipose tissue of subjects was observed after 20 weeks of exposure to the HFHSc diet in comparison to the C group, these results extend those of Peris-Sampedro et al. (2019), who reported an increase of subcutaneous, perirenal and retroperitoneal white adipose tissue. An increase in white adipose tissue had also been observed when subjects were exposed to other types of hypercaloric diets, *e.g*., in a study carried out by (Poret et al., 2018), the weight of epididymal and retroperitoneal fat depots were larger in subjects fed with a high-fat diet; similar results were obtained by Décarie-Spain et al. (2018) using a saturated-high fat diet.

When glucose levels were evaluated after 6 h of fasting, subjects exposed to HFHSc diet presented higher glucose levels in comparison to those obtained in the C group. This result is at odds with those observed by Peris-Sampedro et al. (2019), who found no differences between the group fed with standard chow and the group fed with the choice diet for 4 weeks. This difference could be associated to the length of exposure to the diet, suggesting that four weeks of diet may not have been enough to produce changes in glucose metabolism. In fact, Ebrahim et al. (2021) reported that subjects had higher serum levels of glucose after 16 weeks of a high-fat diet, supporting the hypothesis that changes in glucose levels can be observed when extended periods of exposure to hypercaloric diets are used.

To evaluate the alterations in motivation, we trained subjects in a PR schedule and compared their performance before and during the diet-induced obesity protocol. We found that subjects of both the C and E groups decreased their breaking point after the access to their respective diets; a larger decrease was observed after 2 weeks of exposure to the HFHSc diet in the E group, compared to the C group. The C group showed a decrease in its BP similar to that observed in the E group after 3 weeks of standard food diet. In both cases the observed decrease extended up to the end of the evaluation.

The observed decrease of BP in the E group after the HFHSc diet exposure is in accordance with the report by Tracy et al. (2015), who observed that as the exposure period to the diet lengthens, the motivation to obtain a reinforcer decreased. However, a question emerges: if there is a decrease in the motivation to obtain palatable food, how is it possible that subjects continue consuming the HFHSc diet but still gain body weight? To attempt an explanation to such question, the ways to assess motivation should be addressed. It has been suggested that motivated behavior has activational (promotes an action) and directional components (direct the behavior toward or away from a particular stimulus) (Correa et al., 2020). The activational component is driven by mesolimbic dopamine and supports instrumental responding (Salamone et al., 2022). When dopamine antagonists such as haloperidol were administered, subjects decreased the number of lever presses in a FR5 schedule to obtain palatable pellets and concomitantly, it increased the intake of free-chow when both options were given simultaneously (Salamone et al., 1991). Based on such results it was suggested that mesolimbic dopamine modify the sensitivity of the subjects to the work requirements of operant schedules (Salamone et al., 2022). As mesolimbic dopamine function is altered after subjects are exposed to a saturated high-fat diet (Hryhorczuk et al., 2016), it is possible that subjects could be more sensitive to the response requirements of the PR schedule and are unwilling to work for the reinforcer but continue consuming the palatable food when provided without a response requirement. Additional support for this suggestion is the observation that in the initial evaluation of PR during HFHSc diet access, the PR in the E group was lower than in the C group.

Regarding the experimental design, we evaluated the performance of both groups on the PR schedule, first under food deprived conditions and then under satiety conditions. The difference between these two conditions could explain the decrease observed in the two groups, but not the lack of differences between groups during the diet-induced obesity protocol, nor the fast decrease in BP observed in the E group compared to the C group. Therefore, it is important to consider the conditions of the PR schedule performance. For example, Tracy et al. (2015) reported significant differences between control and experimental groups after a high-fat diet exposure, but their subjects were evaluated in food deprived conditions. Since the PR schedule is sensitive to changes in body weight and satiety conditions (Olarte-Sánchez et al., 2015) is possible that *ad libitum* access to food, independent of the diet, could mask the effects of HFHSc diet on motivation. It is worth noticing that La Fleur et al. (2007) also observed differences between groups under satiety conditions in a PR schedule, these discrepancies with our results could be attributed to the time of exposure to the diet; in their study, exposure to the diet was shorter than ours: 14 days before operant conditioning training.

Another factor to consider is the pre-exposure to the reinforcer before the evaluation, subjects in this experiment received 0.02 ml of a 10% sugar solution as a reinforcer in the PR schedule; this was the same concentration of sucrose solution that was offered in the HFHSc diet. It is possible that the previous experience of sucrose (free access to the solution) affected the performance of the subjects in the PR schedule during the HFHSc diet. In accordance to this suggestion, Vendruscolo et al. (2010) observed that if rats had a previous experience with sucrose during adolescence, they achieved lower BPs during adulthood. In addition, Tracy et al. (2015), also pre-exposed subjects fed with chow and subjects fed with a high-fat diet to the reinforcer and then, compared the BP achieved by both groups and found no differences between groups.

A limitation of our study is that the base line of the BP was obtained under food deprived conditions and the reinforcer was similar to the solution offered on the hypercaloric diet; for future research it would be worth registering the BP under satiety conditions receiving standard diet and using of sucrose pellets as reinforcers. As Salamone (2006) suggested, in a PR schedule subjects also take decisions based in the effort required to obtain the reinforcer; therefore, it may be worth to explore the use of the effort-based decision making task paradigm (Salamone et al., 2016a). In addition, it should be noted that although the BP is the most common parameter obtained in the PR schedule and is widely accepted as an index of motivation, it is also sensitive to non-motivational factors such as the ratio step size, therefore, it would be useful to use alternative analysis of the PR performance (Bradshaw and Killeen, 2012).

Taken together, our results suggest that exposure to a HFHSc diet produced a significant decrease in motivation after 2 weeks that is similar to that observed after 20 weeks of exposure to the diet. The decrease in motivation took longer (3 weeks) to develop using chow diet in the C group. This finding could be of practical use for the design of future investigations of dietary induction of pathological obesity. Furthermore, the use of a HFHSc diet is a useful animal model to study motivational changes during the development of obesity, as exposure to this det produced pathophysiological and psychological alterations observed with other types of hypercaloric diet.

## References

Arcego, D.M., Krolow, R., Lampert, C., Toniazzo, A.P., Garcia, E. dos S., Lazzaretti, C., Costa, G., Scorza, C., Dalmaz, C., 2020. Chronic high-fat diet affects food-motivated behavior and hedonic systems in the nucleus accumbens of male rats. Appetite 153, 104739. https://doi.org/10.1016/j.appet.2020.104739

Berridge, K.C., Ho, C.Y., Richard, J.M., Difeliceantonio, A.G., 2010. The tempted brain eats: Pleasure and desire circuits in obesity and eating disorders. Brain Res. 1350, 43–64. https://doi.org/10.1016/j.brainres.2010.04.003

Bradshaw, C.M., Killeen, P.R., 2012. A theory of behaviour on progressive ratio schedules, with applications in behavioural pharmacology. Psychopharmacology (Berl). 222, 549–564. https://doi.org/10.1007/s00213-012-2771-4

Correa, M., Pardo, M., Carratalá-Ros, C., Martínez-Verdú, A., Salamone, J.D., 2020. Preference for vigorous exercise versus sedentary sucrose drinking: An animal model of anergia induced by dopamine receptor antagonism. Behav. Pharmacol. 31, 553–564. https://doi.org/10.1097/FBP.0000000000000556

Décarie-Spain, L., Sharma, S., Hryhorczuk, C., Issa-Garcia, V., Barker, P.A., Arbour, N., Alquier, T., Fulton, S., 2018. Nucleus accumbens inflammation mediates anxiodepressive behavior and compulsive sucrose seeking elicited by saturated dietary fat. Mol. Metab. 10, 1–13. https://doi.org/10.1016/j.molmet.2018.01.018

Fam, J., Clemens, K.J., Westbrook, R.F., Morris, M.J., Kendig, M.D., 2022. Chronic exposure to cafeteria-style diet in rats alters sweet taste preference and reduces motivation for, but not ‘liking’ of sucrose. Appetite 168. https://doi.org/10.1016/j.appet.2021.105742

Figlewicz, D.P., Jay, J.L., Acheson, M.A., Magrisso, I.J., West, H., Zavosh, A., Benoit, S.C., Davis, J.F., 2014. NIH Public Access 61, 19–29. https://doi.org/10.1016/j.appet.2012.09.021.Moderate

Furman, B.L., 2015. Streptozotocin-Induced Diabetic Models in Mice and Rats. Curr. Protoc. Pharmacol. 70, 5.47.1–5.47.20. https://doi.org/10.1002/0471141755.ph0547s70

Gao, Y., Bielohuby, M., Fleming, T., Grabner, G.F., Foppen, E., Bernhard, W., Guzmán-Ruiz, M., Layritz, C., Legutko, B., Zinser, E., García-Cáceres, C., Buijs, R.M., Woods, S.C., Kalsbeek, A., Seeley, R.J., Nawroth, P.P., Bidlingmaier, M., Tschöp, M.H., Yi, C.X., 2017. Dietary sugars, not lipids, drive hypothalamic inflammation. Mol. Metab. 6, 897–908. https://doi.org/10.1016/j.molmet.2017.06.008

Hryhorczuk, C., Florea, M., Rodaros, D., Poirier, I., Daneault, C., Des Rosiers, C., Arvanitogiannis, A., Alquier, T., Fulton, S., 2016. Dampened mesolimbic dopamine function and signaling by saturated but not monounsaturated dietary lipids. Neuropsychopharmacology 41, 811–821. https://doi.org/10.1038/npp.2015.207

La Fleur, S.E., Vanderschuren, L.J.M.J., Luijendijk, M.C., Kloeze, B.M., Tiesjema, B., Adan, R.A.H., 2007. A reciprocal interaction between food-motivated behavior and diet-induced obesity. Int. J. Obes. 31, 1286–1294. https://doi.org/10.1038/sj.ijo.0803570

Lutz, T.A., Woods, S.C., 2012. Overview of animal models of obesity. Curr. Protoc. Pharmacol. 1–22. https://doi.org/10.1002/0471141755.ph0561s58

Olarte-Sánchez, C.M., Valencia-Torres, L., Cassaday, H.J., Bradshaw, C.M., Szabadi, E., 2015. Quantitative analysis of performance on a progressive-ratio schedule: Effects of reinforcer type, food deprivation and acute treatment with δ9-tetrahydrocannabinol (THC). Behav. Processes 113, 122–131. https://doi.org/10.1016/j.beproc.2015.01.014

Ortiz, V.E., Kwo, J., 2015. Obesity: Physiologic changes and implications for preoperative management. BMC Anesthesiol. 15, 1–12. https://doi.org/10.1186/s12871-015-0079-8

Peris-Sampedro, F., Mounib, M., Schéle, E., Edvardsson, C.E., Stoltenborg, I., Adan, R.A.H., Dickson, S.L., 2019. Impact of Free-Choice Diets High in Fat and Different Sugars on Metabolic Outcome and Anxiety-Like Behavior in Rats. Obesity 27, 409–419. https://doi.org/10.1002/oby.22381

Poret, J.M., Souza-Smith, F., Marcell, S.J., Gaudet, D.A., Tzeng, T.H., Braymer, H.D., Harrison-Bernard, L.M., Primeaux, S.D., 2018. High fat diet consumption differentially affects adipose tissue inflammation and adipocyte size in obesity-prone and obesity-resistant rats. Int. J. Obes. 42, 535–541. https://doi.org/10.1038/ijo.2017.280

Roberts, D.C.S., Richardson, N.R., 2003. Self-Administration of Psychomotor Stimulants Using Progressive Ratio Schedules of Reinforcement. Anim. Model. Drug Addict. 24, 233–270. https://doi.org/10.1385/0-89603-217-5:233

Salamone, J.D., 2006. Will the last person who uses the term “reward” please turn out the lights? Comments on processes related to reinforcement, learning, motivation and effort. Addict. Biol. 11, 43–44. https://doi.org/10.1111/j.1369-1600.2006.00011.x

Salamone, J.D., Correa, M., Yohn, S., Lopez Cruz, L., San Miguel, N., Alatorre, L., 2016a. The pharmacology of effort-related choice behavior: Dopamine, depression, and individual differences. Behav. Processes 127, 3–17. https://doi.org/10.1016/j.beproc.2016.02.008

Salamone, J.D., Ecevitoglu, A., Carratala-Ros, C., Presby, R.E., Edelstein, G.A., Fleeher, R., Rotolo, R.A., Meka, N., Srinath, S., Masthay, J.C., Correa, M., 2022. Complexities and paradoxes in understanding the role of dopamine in incentive motivation and instrumental action: Exertion of effort vs. anhedonia. Brain Res. Bull. 182, 57–66. https://doi.org/10.1016/j.brainresbull.2022.01.019

Salamone, J.D., Steinpreis, R.E., McCullough, L.D., Smith, P., Grebel, D., Mahan, K., 1991. Haloperidol and nucleus accumbens dopamine depletion suppress lever pressing for food but increase free food consumption in a novel food choice procedure. Psychopharmacology (Berl). 104, 515–521. https://doi.org/10.1007/BF02245659

Salamone, J.D., Yohn, S.E., López-Cruz, L., San Miguel, N., Correa, M., 2016b. Activational and effort-related aspects of motivation: Neural mechanisms and implications for psychopathology. Brain 139, 1325–1347. https://doi.org/10.1093/brain/aww050

Sharma, S., Fulton, S., 2013. Diet-induced obesity promotes depressive-like behaviour that is associated with neural adaptations in brain reward circuitry. Int. J. Obes. 37, 382–389. https://doi.org/10.1038/ijo.2012.48

Smith, K.B., Smith, M.S., 2016. Obesity Statistics. Prim. Care - Clin. Off. Pract. 43, 121–135. https://doi.org/10.1016/j.pop.2015.10.001

Tracy, A.L., Wee, C.J.M., Hazeltine, G.E., Carter, R.A., 2015. Characterization of attenuated food motivation in high-fat diet-induced obesity: Critical roles for time on diet and reinforcer familiarity. Physiol. Behav. 141, 69–77. https://doi.org/10.1016/j.physbeh.2015.01.008

Valencia-Torres, L., Bradshaw, C.M., Bouzas, A., Hong, E., Orduña, V., 2014. Effect of streptozotocin-induced diabetes on performance on a progressive ratio schedule. Psychopharmacology (Berl). 231, 2375–2384. https://doi.org/10.1007/s00213-013-3401-5

Vendruscolo, L.F., Gueye, A.B., Darnaudéry, M., Ahmed, S.H., Cador, M., 2010. Sugar overconsumption during adolescence selectively alters motivation and reward function in adult rats. PLoS One 5. https://doi.org/10.1371/journal.pone.0009296

